# Acetyl-CoA production by select metabolic pathways promotes cardiac repair after myocardial infarction via mediating histone acetylation

**DOI:** 10.1101/622746

**Authors:** Ienglam Lei, Shuo Tian, Wenbin Gao, Liu Liu, Yijing Guo, Zhong Wang

## Abstract

Myocardial infarction (MI) is accompanied by severe energy deprivation and extensive epigenetic changes. However, how energy metabolism and chromatin modifications are interlinked during MI and heart repair has been poorly explored. Here, we examined the effect of different carbon sources that involved in the major metabolic pathways of acetyl-CoA synthesis on myocardial infarction and found that elevation of acetyl-CoA significantly improved heart function in I/R rats by administration of sodium octanoate (8C). Mechanistically, 8C prevented I/R injury by promoting histone acetylation which in turn activated the expression of antioxidant genes HO1, NQO1 and SOD2 and inhibited cardiomyocyte apoptosis. Furthermore, we identified that 8C-promoted histone acetylation and heart repair were carried out by metabolic enzyme medium-chain acyl-CoA dehydrogenase (MCAD) and histone acetyltransferase Kat2a. Therefore, our results demonstrate that 8C dramatically improves cardiac function through metabolic acetyl-CoA-mediated histone acetylation. This study uncovers an interlinked metabolic/epigenetic network comprising 8C, acetyl-CoA, MCAD, and Kat2a in stimulating histone acetylation and anti-oxidative stress gene expression to combat heart injury.

## Introduction

Energy is a fundamental requirement for all living organisms and its production typically requires fuels (metabolites) and oxygen. Heart is a most sensitive organ to energy supply and provision. One key response to fuel changes is the epigenetic modifications using one carbon or two carbon moieties derived from metabolites to change chromatin structure by methylation and acetylation and regulate gene expression^1^. Notably, a bi-directional interplay between metabolism and epigenetic control has been proposed recently: metabolism directly regulates chromatin epigenetic state, whereas chromatin state defines gene control in response to metabolic status^2,3^.

Myocardial infarction (MI), which blocks the supply of energy to infarcted area, is one of the leading causes of death in the world^4^. Despite the severe complications of this devastating disease^5^, the therapies to reduce MI injury are still limited. MI is accompanied by severe energy deprivation and extensive epigenetic changes. Regulations in either energy metabolism or epigenetics are essential for heart function and pathogenesis^6,7^, but how these two events are interlinked in the context of MI has been under-explored. Clearly, exploring an integrated metabolic and chromatin control in MI injury and heart repair and regeneration may provide novel therapies against heart injury.

Acetyl-CoA, a key metabolite of energy-providing substrate, also serve as the substrate for histone acetyltransferases (HATs) to transfer the acetyl-group to histone residues for histone acetylation^8,9^. Manipulation of acetyl-CoA, either by intervention of synthetic enzymes, or nutrient source could alter histone acetylation in various cell types^9,12^. Importantly, acetyl-CoA induces metabolic adaptations through regulating histone acetylation in response to starving or hypoxia conditions^13,14^. Moreover, myocardial ischemia reperfusion (I/R) injury causes dramatic metabolism changes and subsequent histone deacetylation. Inhibition of histone deacetylation activity protects heart function after I/R injury^15–17^ and inhibits cardiac remodeling and heart failure^18^. Thus, the intersection of acetyl-CoA-mediated metabolism and histone acetylation is very likely a novel hub for identifying targets for heart repair.

In this study, we examined whether metabolic manipulation of acetyl-CoA level could alter histone acetylation, which in turn promote heart repair and protection after I/R injury. I/R injury was accompanied with dramatic decreases in acetyl-CoA and histone acetylation. Our screen identified a nutrient, the medium chain fatty acid 8C, as an effective carbon source to stimulate acetyl-CoA production and histone acetylation and protect heart function after MI. Specifically, we showed that a single i.p. injection of 8C at the time of reperfusion significantly reduced the infarct size and improved cardiac function at both 24 hours and 4 weeks after MI. We found that 8C produced acetyl-CoA and rescued histone acetylation decrease both in vivo and in vitro after I/R injury, which led to elevated expression of antioxidant genes. Importantly, we elucidated that the metabolic enzyme medium-chain acyl-CoA dehydrogenase (MCAD) and histone acetyltransferase Kat2a were key factors in transferring the acetyl moiety from 8C to acetyl groups in cardiomyocyte histone acetylation for I/R injury repair. We further determined that a major function of 8C, MCAD, and Kat2a mediated histone acetylation in heart repair was to stimulate gene expression to alleviate oxidative stress in cardiomyocytes. Thus, our study established a novel interlinked metabolic/epigenetic network that may provide new strategies to treat heart injuries.

## Results

### Administration of 8C at reperfusion elevated acetyl-CoA and improved short-term and long-term cardiac function after I/R

Myocardial infarction (MI) induces dramatic metabolic and epigenetic changes including decrease of acetyl-CoA synthesis and histone acetylation^19,20^. We reasoned that swift synthesis of acetyl-CoA could rescue heart function by restoring histone acetylation. Based on the major metabolic pathways for acetyl-CoA synthesis (Figure 1A) ^2^, we examined the effect of sodium acetate (2C, 500 mg/kg), sodium pyruvate (3C, 500 mg/kg), sodium citrate (5C, 500 mg/kg), sodium octanoate (8C, 160 mg/kg), and sodium nonanoate (9C, 200 mg/kg) on heart function after I/R. Rats were intraperitoneally (i.p.) injected with these metabolites as well as saline for three days and another dose before I/R surgery (Figure 1B). At 24 hours after I/R, the infarct size was measured by Evans blue and tripheyltetrazolium chloride (TTC) staining at 24 hours after I/R. Evans blue staining indicated area at risk (AAR) and TTC staining indicated the infarct size (IS). The ratio of AAR/LV was similar in all groups (Figure S1A-B). Interestingly, these metabolites displayed distinct effects on reducing myocardial infarct size. Among the five metabolites examined, 2C, 3C, and 8C significantly reduced the IS/AAR ratio, whereas 5C and 9C treatment did not reduce IS/AAR ratio compared to saline treated groups (Figure 1C and Figure S1A). These results indicated that interventions of acetyl-CoA by modulating select metabolic pathways could improve cardiac repair after I/R.

**Figure 1.**
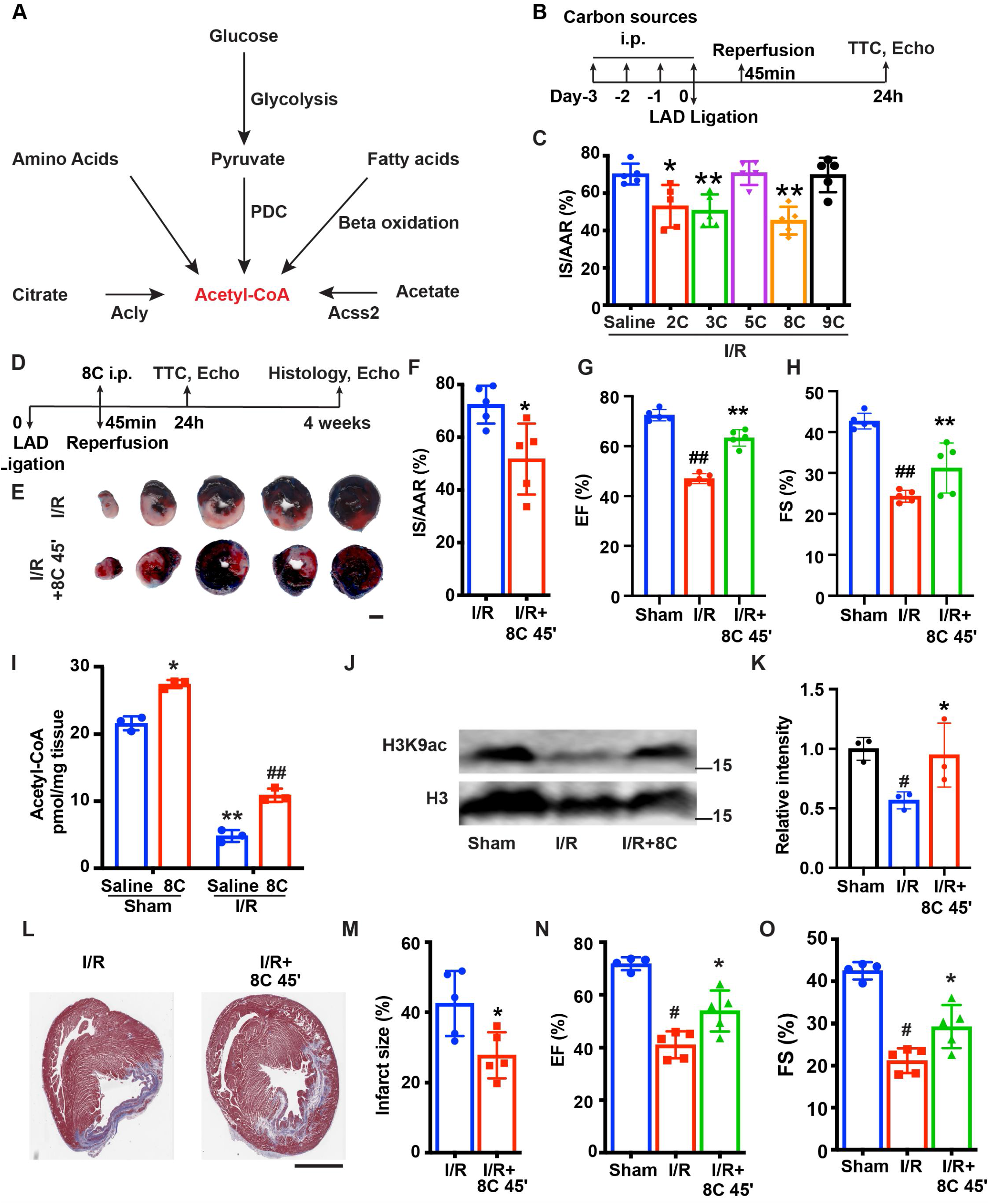
A screen of acetyl-CoA synthesis pathways identified 8C as a potent reagent to protect against ischemia-reperfusion injury in rats. (A) Metabolic pathways of acetyl-CoA synthesis. (B) Schematic diagram of different carbon sources administration prior to I/R surgery. (C) Quantification of IS/AAR ratio in Figure S1A by Image J. (D) Schematic diagram of 8C administration post ischemic injury. (E) Representative figures of heart sections at 24 hours after I/R with or without 8C administration at reperfusion after 45 minutes ischemia. Scale bar: 2.5 mm. (F) Quantification of IS/AAR ratio in Figure 1E. LV EF (G) and FS (H) at 24 hours after I/R. (I) Quantification of Acetyl-CoA levels in sham and I/R rat hearts at indicated conditions. (J-K) Western blot and quantification of H3K9ac and H3 in rat hearts 24 hours after I/R. (L) Trichrome masson staining of heart section after 4 weeks of I/R. Scale bar: 2.5 mm.(M) Quantification of infarct size in Figure 1I. LV EF (N) and FS (O)at 4 weeks after I/R. Error bars represent S.D. n=5, ^#^p<0.05, ^##^p<0.01 vs Sham group. * p<0.05, **p<0.01, vs I/R group.

As 8C administration resulted in the most dramatic protection among all the metabolites tested after I/R (Figure 1C), we focused on investigating the role of 8C on heart protection in a more clinically relevant setting. 8C (160 mg/kg) or saline were i.p. injected only once at the time of reperfusion, which was 45 minutes after LAD ligation. The infarct size was then measured by Evans blue/TTC staining at 24 hours after I/R (Figure 1D). Evans blue/TTC staining showed that 24 hours after reperfusion, 8C administration reduced the IS/AAR ratio from 72% to 52% compared to saline control after I/R (Figure 1E-F and Figure S1C). Moreover, 8C also significantly improved the left ventricle function evaluated by echocardiography 24 hours after I/R. Compared to saline treated I/R rats, 8C administration led to increase of EF and FS from 47% to 58% and 24% to 31%, respectively (Figure 1G-H), while the normal rat heart has EF at 72% and FS at 43%. These results indicated that 8C significantly improve cardiac function by approximately 40%. Importantly, we found that there was a significant reduction of acetyl-CoA concentration in hearts after I/R, while 8C administration increased acetyl-CoA level in hearts after I/R (Figure 1I). Moreover, 8C administration elevated H3K9ac level in hearts after I/R (Figure 1J-K). Furthermore, to investigate whether a single dose of 8C administration is beneficial for long-term cardiac function after I/R, we examined the infarct size and heart function at 4 weeks after I/R. Trichrome Masson staining showed that the infarct size was notably reduced after 8C treatment (Figure 1L-1M). The left ventricle function was also improved after 8C treatment as evidenced by the increase of EF and FS (Figure 1N-1O). Together with the fact that 8C can quickly enter into cells and contribute to around 50% of acetyl-CoA in heart at one hour after administration^21^, our results indicated that 8C administration at the time of reperfusion elevated acetyl-CoA production and histone acetylation level and significantly improved both short-term and long-term cardiac function after I/R.

### 8C attenuated cardiomyocyte apoptosis through alleviating oxidative stress

Apoptosis is one of the major reasons for cardiac damage after I/R injury^22^. To detect the impact of 8C on cardiomyocyte apoptosis, TUNEL staining was performed at the border zone of infarct hearts. 8C treatment dramatically reduced TUNEL positive cardiomyocytes in border zone (Figure 2A-2B). Consistent with the TUNEL assay, the levels of cell death indicators serum CK and LDH were also reduced in I/R rats after 8C administration (Figure 2C-2D). Moreover, 8C led to reduction in pro-apoptotic regulator Bax and upregulation of anti-apoptotic gene Bcl2 at 24 hours after I/R injury (Figure 2E-2F). To study the mechanism of the beneficial effect of 8C after I/R, we examined the gene expression profile at border zone 24 hours after I/R by RNA-seq. GSEA^23^ showed that genes involved in apoptotic signaling pathways were enriched in saline-treated compared to 8C-treated rats after I/R (Figure 2G). GO analysis using differential expressed genes in 8C and saline treated rats showed that genes related to cellular response to tumor necrosis factor were highly enriched in saline treated I/R groups, whereas genes related to cardiomyocyte functions were highly expressed in 8C treated I/R hearts (Figure S2A-B). These results suggested 8C administration reduced the cardiomyocyte death after I/R. Since I/R induced oxidative stress triggers cardiomyocyte apoptosis after reperfusion^24^, it is likely that 8C reduced cell death by activating antioxidant process after I/R injury. To address this, we measured the level of cardiac reactive oxygen species (ROS) after I/R by staining of dihydroethidium (DHE), a chemical that could be oxidized by ROS^25^. The intensity of DHE signal was significantly lower in the presence of 8C after I/R (Figure 2H-2I), indicating that 8C reduced oxidative stress after I/R. Moreover, 8C rescued the myocardial SOD activity after I/R (Figure S2C). Altogether, these results showed that one major mechanism through which 8C improved cardiac function after I/R was reduction of the oxidative stress and subsequent cell apoptosis.

**Figure 2.**
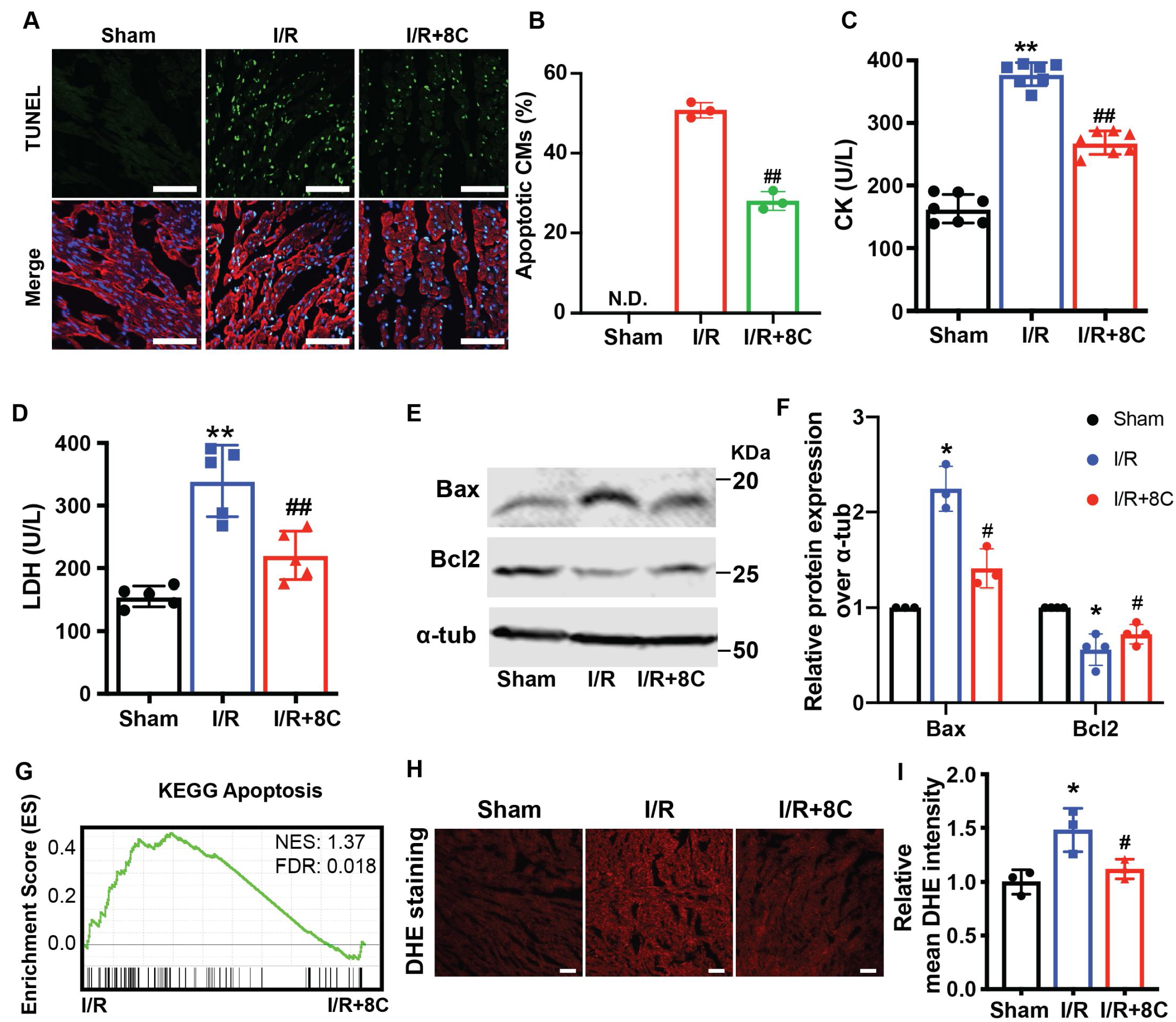
Post ischemic administration of 8C reduced the oxidative stress and cell death after I/R injury. (A) Representative of TUNEL (green) and cTnT (red) double staining at boarder zone at 24 hours post I/R injury. Scale bar: 100 μm (B) Quantification of cardiomyocytes cell death of 12 sections (C-D) Serum CK and LDH level at 24 hours post I/R. (E-F) Western blot and quantification of Bax and 8Cl2 at 24 hours after I/R. (G) GSEA analysis of Kegg apoptotic pathways after I/R with and without 8C treatment. (H) Heatmap of antioxidant genes expression. (I) ROS levels were measured by OHE staining. Scale bar: 200μm. (J) Relative mean OHE fluorescence intensity measured by Image J. Error bars represent S.D. n=4. *p<0.05, **p<0.01 vs Sham; ^#^p<0.05, ^##^p<0.01 vs I/R group.

To further investigate the effect of 8C on oxidative stress and apoptosis in cardiomyocytes, neonatal rat ventricle myocytes (NRVM) were subjected to simulate ischemia reperfusion (sI/R) with or without 8C. NRVM were pretreated with 8C or PBS for 12 hours, and then subjected to 2 hours of simulated ischemia followed by 4 hours of simulated reperfusion. The percentage of apoptotic cells after sI/R was measured by labeling with Annexin V and PI. 8C significantly reduced the percentage of apoptotic cells, as evidenced by the lower percentage of Annexin V and PI double positive cells (Figure 3A-3B). In addition, combination of CCK8 and LDH release assays were used to measure the total cell death^26^. 8C administration reduced cell death after sI/R as showed by increased cell viability and decreased LDH release into to culture medium (Figure 3C-3D). Moreover, 8C reduced the expression of cleaved Caspase 3 after sI/R. Consequently, the expression of cleaved PARP, which is cleaved by Caspase 3 was also reduced with 8C administration (Figure 3E). Altogether, these results demonstrated that 8C reduced the apoptosis of NRVM exposed to sI/R. To assess whether 8C reduced the oxidative stress in cardiomyocytes after sI/R, the accumulation of ROS level was measured by the intensity of DHE staining. The ROS levels were significantly decreased in NRVM with 8C treatment after sI/R (Figure 3F-3H), indicating a direct effect of 8C in alleviating oxidative stress after sI/R. Thus, our collective in vivo and in vitro results revealed that 8C attenuated cardiomyocyte apoptosis through alleviating oxidative stress.

**Figure 3.**
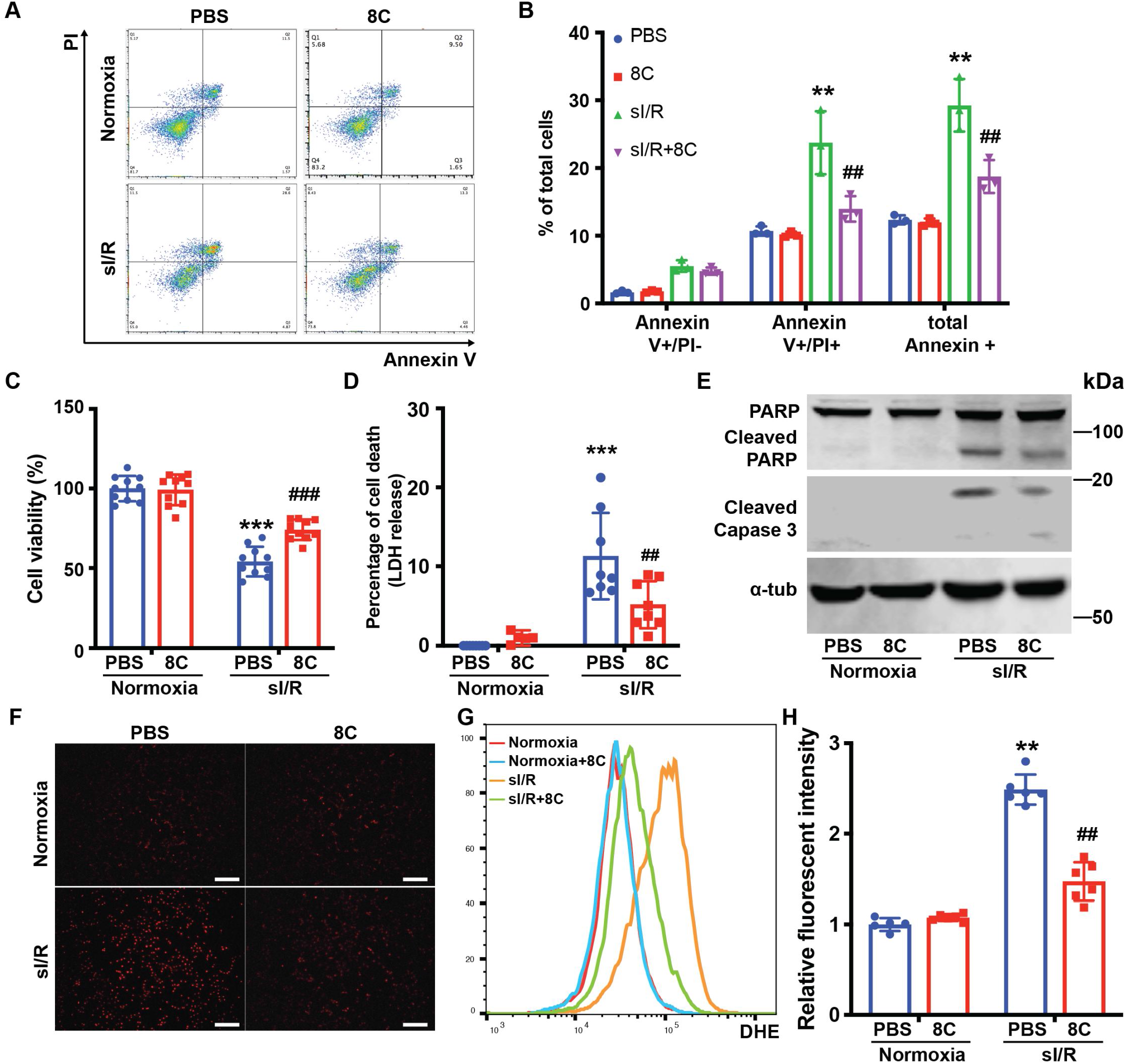
8C attenuated NRVM apoptosis through reducing oxidative stress. (A) FACS analysis of Annexin V and PI staining in NRVM exposed to sI/R with and without 8C treatment. (B) Quantification of percentage of Annexin V+ and PI + cells. Cell viability and cell death measurement in NRVM with sI/R using CCK8 detection kit (C) and LDH assay kit (D). (E) Western blot of cleaved Caspase 3 and cleaved PARP in NRVM after sI/R treatment. (F) NRVM cellular ROS levels were indicated by OHE staining after sI/R treatment. Scale bar: 200μm (G) FACS analysis of DHE staining NRVM after sI/R. (H) Relative mean fluorescence intensity of OHE staining measured by Flowjo. n=3, **p<0.01, ***p<0.001, vs Normo ia+PBS; ^##^ p<0.01,^###^p<0.001 vs sI/R+PBS.

### Increasing acetyl-CoA synthesis by 8C administration stimulated histone acetylation and promoted anti-oxidative gene expression to combat ROS after I/R injury

8C-generated acetyl-CoA contributes to histone acetylation in several cell types^12^, and histone acetylation plays an important role in regulating cellular response to oxidative stress^27^. To examine whether 8C protected cardiomyocytes against I/R injury through stimulating histone acetylation, we first measured the acetyl-CoA level in NRVMs in vitro under sI/R. Similar to in vivo I/R injury, sI/R reduced acetyl-CoA level in NRVMs and 8C significantly increased the production of acetyl-CoA in NRVMs (Figure 4A). To determine the effect of acetyl-CoA replenishment on histone acetylation, we measured H3K9ac, H3K14ac, H3K27ac and total H3 acetylation in NRVM after sI/R. We found that sI/R led to a remarkable decrease of H3K9ac, H3K14ac, H3K27ac and acH3, and that 8C increased histone acetylation histone acetylation in normal NRVM and rescued sI/R reduced acetylation (Figure 4B-E). These results indicated that acetyl-CoA production by 8C rescued histone acetylation decrease after sI/R.

**Figure 4.**
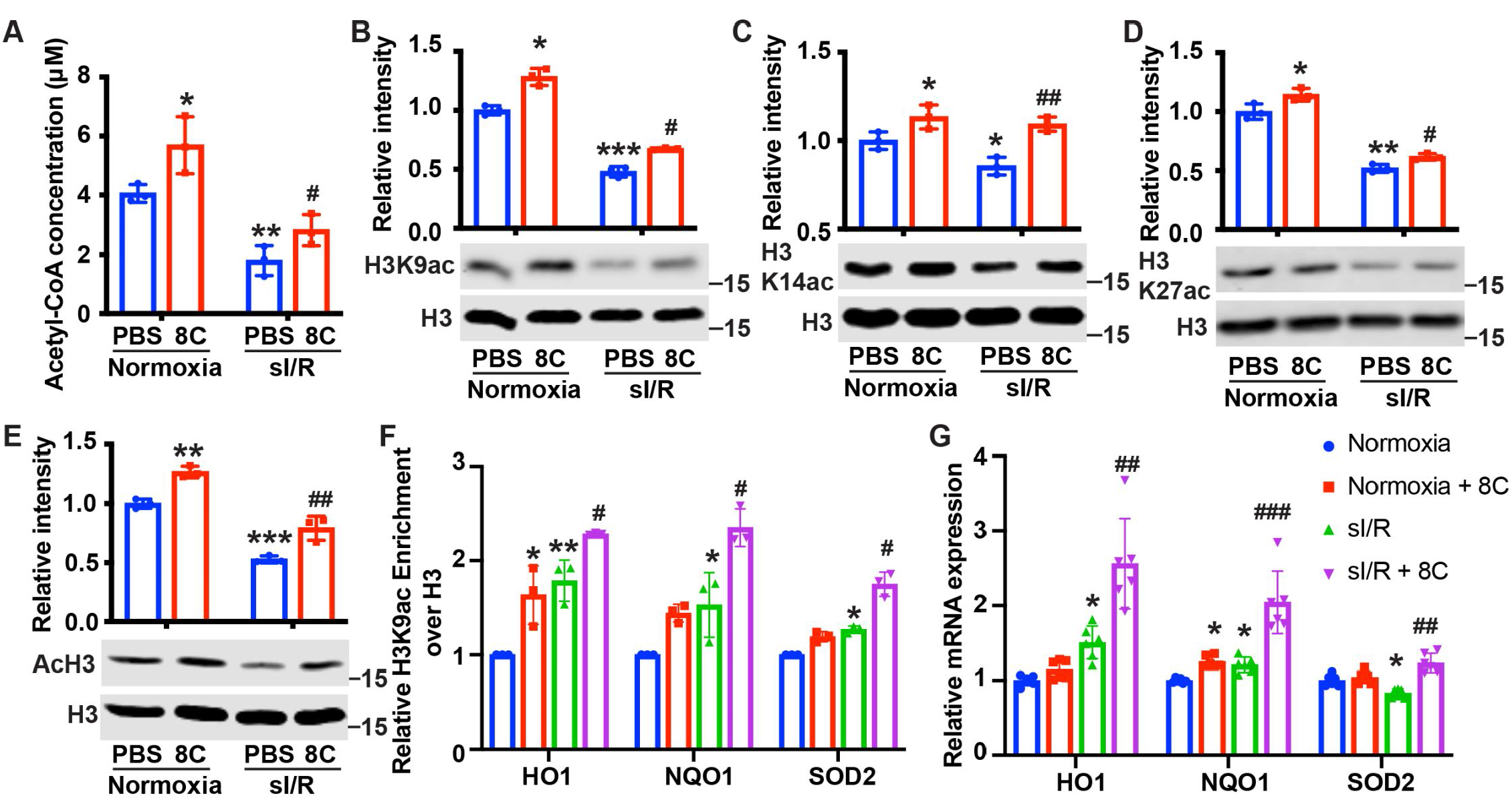
8C stimulated histone acetylation and promotes antioxidant gene expression. (A) Quantification of Acetyl-CoA concentrations in NRVM subjected to sI/R. (B-E) 8C rescues sI/R reduced H3K9ac, H3K27ac, H3K14ac and total acH3 levels. NRVMs were treated with or without 0.5mM 8C under sI/R. The histone acetylation levels were determined by western blot. Total H3 in the same blot was used as loading control. (F) Enrichment of H3K9ac over H3 at promoters of H01, NQ01, and S002 in NRVM after sI/R. (G) mRNA expression of H01, NQ01, and S002 in NRVM after sI/R. *p<0.05, **p<0.01, ***p<0.001, vs Normoxia + PBS; ^#^p<0.05, ^##^p<0.01, ^###^p<0.001 vs sI/R + PBS.

Acetyl-CoA is the substrate for histone acetyltransferases (HATs) to generate histone acetylation by transferring the acetyl-group from acetyl-CoA to histone lysine residues^28^. Specifically, HATs with low affinity to acetyl-CoA are more sensitive to acetyl-CoA abundance. H3K9ac has been reported as the histone acetylation most sensitive to acetyl-CoA levels^28^. Consistent with this finding, we found that 8C led to most significant changes in H3K9ac after sI/R. Thus, we reasoned that H3K9ac, which is enriched in promoters for gene activation, is one key epigenetic event for gene regulation after sI/R. To examine the potential epigenetic regulation of 8C derived acetyl-CoA in anti-oxidative stress, we performed ChIP to measure H3K9ac at the promoters of antioxidant genes. While sI/R led to increased H3K9ac level at the promoters of NQO1, HO1 and SOD2, 8C further elevated H3K9ac on the promoters of these genes (Figure 4F). Consequently, 8C upregulated both mRNA and protein expression of antioxidant genes including HO1, NQO1, and SOD2 after sI/R (Figure 4G and Figure S3). Collectively, our results suggested that 8C-produced acetyl-CoA contributed to epigenetic regulation of antioxidant genes to combat ROS in response to I/R injury.

### MCAD was required for the metabolism of 8C into acetyl-CoA and subsequent histone acetylation increase and heart protection

To ascertain whether 8C produced acetyl-CoA was important for the rescue of histone acetylation after sI/R, we knocked down MCAD (Figure S4A), a key enzyme in the generation of acetyl-CoA from 8C^29^. Knockdown of MCAD disrupted the metabolism of 8C, and therefore led to reduction of histone acetylation promoted by 8C in NRVM in both normoxia and sI/R condition (Figure 5A). Thus, these data indicated that metabolic production of acetyl-CoA from 8C was required for the 8C-mediated histone acetylation regulation. To determine whether acetyl-CoA mediated histone acetylation was key to the 8C heart protective effect after I/R, we examined the cardiomyocyte survival after MCAD knockdown with and without 8C after sI/R. MCAD knockdown significantly blocked the protective effect of 8C after sI/R, as evidenced by decreased cell viability and increased LDH release in MCAD knockdown cells in presence of 8C after sI/R compared to knockdown controls with 8C treatment after sI/R (Figure 5B and Figure S4B). Importantly, MCAD knockdown blocked the 8C-reduced cellular ROS level after sI/R (Figure 5C-D and Figure S4C). Consistent with inhibited ROS reduction, we found that MCAD knockdown blunted 8C stimulated H3K9ac increase in the promoters of HO1 and NQO1 after sI/R (Figure 5E). Subsequently, MCAD knockdown reduced 8C-elevated expression of HO1 and NQO1 after sI/R (Figure 5F and Figure S4D-F). Thus, these results demonstrated that MCAD-mediated 8C metabolism was essential for histone acetylation and attenuating apoptosis through activating antioxidative process.

**Figure 5.**
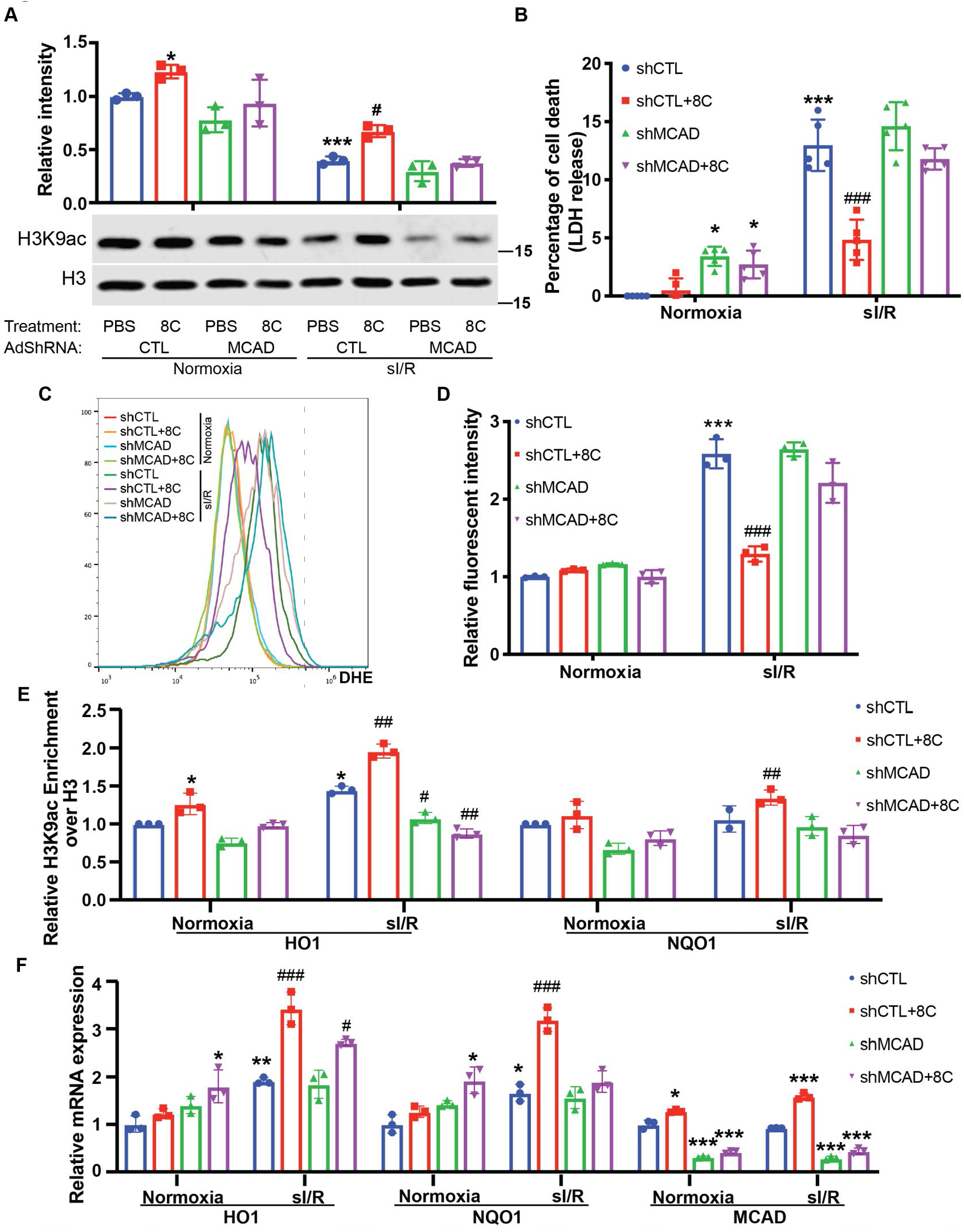
MCAD was required for the conversion of 8C into acetyl-CoA and subsequent histone acetylation increase and heart protection. (A) Western blot of H3K9ac level showed MCAO knockdown reduced 8C-induced H3K9ac increase in NRVM under both normoxia and sI/R. (B) Measurement of medium LDH levels in NRVM at indicated condition using LDH assay kit. (C) FACS analysis of OHE staining NRVM after sI/R. (D) Relative mean fluorescene intensity of DHE staining. (E) Enrichment of H3K9ac over H3 at promoters of HO1 and NQO1 after s|/R at indicated conditions. (F) mRNA expression of HO1, NQO1 and MCAD in NRVM after s|/R. n=3, *p<0.05, **p<0.01, ***p<0.001, vs Normoxia + PBS + shCTL; ^#^p<0.05, ^##^p<0.01, ^###^p<0.001 vs s|/R + PBS + shCTL.

### HAT enzyme Kat2a was required for 8C mediated histone acetylation to inhibit oxidative stress in heart protection

HATs are the very enzymes that catalyze histone acetylation by transferring the acetyl-group from acetyl-CoA to histone lysine residues. As H3K9ac is most sensitive to physiological acetyl-CoA levels, we then hypothesized that HATs that acetylate H3K9 and are most responsive to physiological acetyl-CoA concentrations would play important roles in the cardioprotection of 8C. Kat2a and Kat2b are the major HATs that modulate H3K9ac^30^. Kat2a is mostly responsive to acetyl-CoA concentrations at 0-10 μM^31^, while Kat2b is response to acetyl-CoA in the 0 to 300 μM range^32^. Considering the fact that acetyl-CoA levels in NRVM ranged from 1-7 μM under sI/R (Figure 4A), we focused on studying the effect of Kat2a knockdown on the protective role of 8C after sI/R. Kat2a knockdown largely abolished the H3K9ac increase caused by 8C treatment in NRVM (Figure 6A and Figure S5A), indicating that Kat2a was a key HAT to mediate 8C-stimulated histone acetylation. Furthermore, Kat2a knockdown led to a significant decrease of cell viability and increase of LDH release compared to knockdown control group in presence of 8C after sI/R (Figure 6B and Figure S5B), suggesting that Kat2a was required for the protective effect of 8C. Moreover, knockdown of Kat2a abolished 8C’s effect on inhibiting the cellular ROS level after sI/R (Figure 6C-D and Figure S5C). Specifically, Kat2a knockdown reduced 8C-elevated expression of HO1 and NQO1 after sI/R (Figure 6E and Supplemental Figure S5D-F). Moreover, Kat2a knockdown abolished 8C-stimulated H3K9ac increase at the promoters of HO1 and NQO1 after sI/R (Figure 6F). These results illustrated that Kat2a was required to execute the rescuing role of 8C after sI/R by modulating histone acetylation, which in turn activated antioxidant gene expression and attenuated cellular apoptotic after sI/R. Together our investigation revealed an integrated metabolic and epigenetic network comprising 8C, acetyl-CoA, MCAD, and Kat2a, that likely played an essential role in combating heart injury after I/R (Figure 6G).

**Figure 6.**
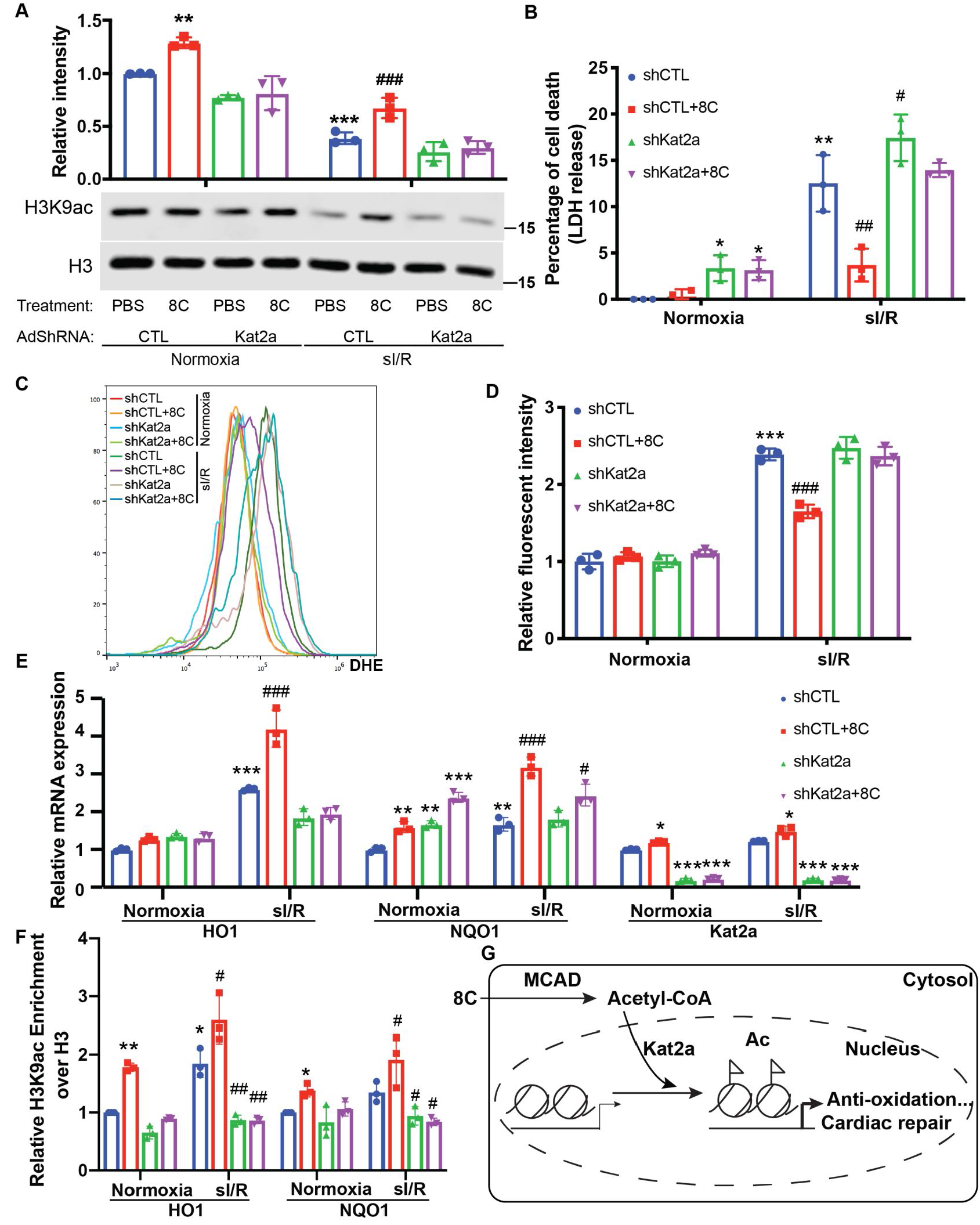
HAT enzyme Kat2a was required for 8C mediated histone acetylation to inhibit oxidative stress in heart protection. (A) Western blot of H3K9ac level showed Kat2a knockdown reduced 8C-induced H3K9ac increase in NRVM under both normoxia and s|/R, (B) Measurement of medium LDH levels in NRVM at indicated condition using LDH assay kit. (C) FACS analysis of DHE staining NRVM after s|/R. (D) Relative mean fluorescene intensity of DHE staining. (E) Enrichment of H3K9ac over H3 at promoters of HO1 and NQO1 after s|/R at indicated conditions. (F) mRNA expression of HO1, NQO1 and Kat2a in NRVM after s|R. n=3, *p<0.05, **p<0.01, ***p<0.001, vs Normoxia+PBS+shCTL; ^#^p<0.05, ^##^p<0.01, ^###^p<0.001 vs s|/R+PBS+shCTL.

## Discussion

In this study, we have established an interlinked metabolic and epigenetic network comprising 8C, acetyl-CoA, MCAD, and Kat2a that stimulates histone acetylation and anti-oxidative stress gene expression to combat heart injury (Figure 6G). Our screen of acetyl-CoA synthesis metabolites identifies that 8C administration significantly improves cardiac function after I/R injury. Specifically, we establish that induction of acetyl-CoA synthesis by 8C metabolism stimulates histone acetylation and promotes cardiomyocyte survival after I/R. Our study further reveals that 8C-stimulated histone acetylation leads to increase of antioxidant gene expression for heart repair. Moreover, MCAD knockdown diminishes the 8C-induced acetylation and subsequently lowers antioxidant activity after sI/R in vitro, indicating that the metabolic conversion of 8C to acetyl-CoA is mainly responsible for histone acetylation and subsequent heart repair effect. Furthermore, the effect of 8C on heart repair through acetyl-CoA and subsequent nuclear histone acetylation is evidenced by Kat2a studies, as Kat2a knockdown largely diminishes the protective effect of 8C after sI/R. Our study demonstrates systematically for the first time that modulating acetyl-CoA abundance can determine cardiomyocyte response to I/R injury via common metabolic and epigenetic mechanisms.

Our study reveals a novel mechanism centered on acetyl-CoA that connects metabolic dynamics and epigenetic regulation in cardiac repair after I/R injury and suggests that acetyl-CoA could be a survival signal for cardiomyocyte after I/R injury. Our data show that I/R injury reduces the cellular acetyl-CoA level, which is associated with decrease of histone acetylation and increase of cardiomyocyte death after I/R injury^20^. We further show that these associations are causally related. In particular, we demonstrate that 8C restores the acetyl-CoA level and subsequently increases the histone acetylation and improves cardiac function after injury. Moreover, 8C administration directly contribute the about 50% of acetyl-CoA in rat hearts^21^, implying that 8C derived acetyl-CoA play an essential role after I/R injury. Importantly, knockdown of 8C metabolic enzyme MCAD, which metabolizes 8C to acetyl-CoA, diminishes the rescuing effect of 8C treatment. Thus, our study suggests that acetyl-CoA from 8C metabolism could improve heart function after I/R.

Our study indicates that histone acetylation is a major downstream event of 8C and acetyl-CoA in heart repair after injury, which is consistent with the notion that acetyl-CoA could serve as second messenger to modulate epigenetic response to environmental changes^2^. Knockdown of Kat2a, a major HAT enzyme in catalyzing histone acetylation, greatly diminishes the cardiomyocyte protective effect of 8C metabolism. Consistent with this notion, 8C effect on elevating histone acetylation is largely abolished under Kat2a knockdown conditions. Histone acetylation as a general epigenetic regulatory mechanism can play essential roles in numerous cellular processes^33^. In this study, we have identified that 8C-mediated histone acetylation decreases the ROS level by promoting the expression of HO1, NQO1 and SOD2 after I/R injury in both in vivo and in vitro. These data indicate that the rescuing effect of 8C treatment after heart injury is at least partially through stimulating gene expression against oxidative stress, which is consistent with the previous observation that a high level of histone acetylation activates gene expression against oxidative stress^27^. Thus, our results suggest metabolism dependent acetyl-CoA synthesis provides an important mechanism for histone acetylation to combat heart injury.

Histone acetylation can also be regulated by histone deacetylase (HDACs)^33^. Interestingly, studies by us and others reveal that chemical inhibition of HDACs also leads to attenuation of myocardial infarction^15,16^ and heart failure^18^. Our recent work indicates that valproic acid, an FDA approved HDAC inhibitor for treatment bipolar disorder, protects heart function after I/R injury by promoting a Foxm1-mediated transcriptional pathway^15^. In addition, SAHA, another HDAC inhibitor, blunts myocardial infarction via regulating autophagy activities^16^. Moreover, HDAC inhibitor could alter the acetylation of myofibrils and govern diastolic function of heart^18^. Whether acetyl-CoA mediated histone acetylation and HDAC inhibition share the same regulatory mechanisms in cardiac repair requires detailed investigation. It will also be interesting to determine how metabolism mediated histone acetylation and HDAC inhibition coordinate their actions in cardiac repair.

Our establishment of direct connection between epigenetic status and metabolite abundance in heart repair may provide an alternative perspective for numerous published studies. For instance, activation of AMPK shows cardioprotective effect after I/R injury^34^. It is postulated that the ability of AMPK in enhancing glucose uptake^34^ and suppressing ribosome biogenesis^35^ is the major reason for cardioprotection. However, the translation of AMPK activation for cardiac therapy has not been successful, partially due to limited understanding of the mechanism for AMPK protection. A recent study shows that activation of AMPK results in increased level of acetyl-CoA and therefore likely elevates histone acetylation^36^. Thus, it is possible that AMPK promotes cardiac repair via direct epigenetic regulation. Similarly, altering the levels of different metabolites show beneficial effect on cardiac repair^37,38^. Considering the potential roles of these metabolites in epigenetic regulations, re-examining these studies will likely provide new insights into manipulating metabolites to alter epigenetics in heart disease treatment, and lead to more successful therapy.

Our study suggests that increasing histone acetylation by acetyl-CoA metabolic pathways is an effective strategy for heart repair. Although increasing histone acetylation by metabolic acetyl-CoA production is an effective strategy for heart repair, not all metabolites that generate acetyl-CoA and histone acetylation have the same beneficial effect. One major reason could be that certain metabolites also generate other signals that cancel or even out-weight this beneficial effect. For example, several metabolites, such as succinate, are found to be the major source of ROS production after I/R, and therefore, succinate administration aggravates I/R injury^39,40^. 9C produces succinate through anaplerotic reaction, and our results show that the accumulation of succinate level is much higher in 9C compared to 8C treatment (Figure S1G). These data may explain the null effect of 9C administration in heart repair. Overall, this study elucidates that exploring detailed metabolic and epigenetic mechanisms mediated by various metabolic carbon sources in combating I/R injury will be an exciting research area to develop potential effective therapies for heart and other organs.

## Methods

### Animal experiment

All experiments were approved by the Institutional Animal Care and Use Committee of the University of Michigan and were performed in accordance with the recommendations of the American Association for the Accreditation of Laboratory Animal Care.

### Generation of rat I/R Models

Myocardial ischemia/reperfusion was carried out in rats as described previously^15^. Briefly, 8-9 weeks old SD rats (Charles River Laboratories) were anaesthetized with ketamine (100 mg/kg) and xylazine (10 mg/kg). Myocardial ischemia was performed by occlusion of the left anterior descending coronary artery (LAD) using 6-0 silk sutures. After 45 min of ischemia, the myocardium was reperfused. Sodium acetate (2C), sodium pyruvate (3C), sodium citrate(5C), sodium octanoate (8C), and sodium nonanoate (9C) were dissolved in saline at concentrations of 125mg/mL, 125mg/mL, 125mg/mL, 40mg/mL, and 50mg/mL, respectively. 2C (500 mg/kg) ^41^, 3C (500 mg/kg) ^42^, 5C (500 mg/kg), 8C (160 mg/kg) ^43^ and 9C (200 mg/kg) ^43^ were intraperitoneal (i.p.) injected to rats for 3 continuous days and another dose before LAD ligation for screening assay. In a clinically relevant setting, 8C was i.p. injected at the time of reperfusion, which was 45 minutes after LAD ligation.

### Echocardiography

Echocardiography (ECG) was performed after surgery. Left ventricular internal diameter end diastole (LVIDd) and left ventricular internal diameter end systole (LVIDs) were measured perpendicularly to the long axis of the ventricle. Ejection fraction (EF) and fractional shortening (FS) were calculated according to LVIDd and LVIDs.

### Evans blue/Triphenyltetrazolium chloride (TTC) staining

After 24 hours reperfusion, the LAD was re-occluded and 5% Evens blue was injected into the right ventricle. The heart was then removed, frozen rapidly and sliced into five 2 mm transverse sections. The sections were incubated with 1% TTC in phosphate buffer (pH 7.4) at 37°C for 10 minutes and photographed by EPSON scanner. The ischemia area (IS), area at risk (AAR), and left ventricular area, were measured with Image J software.

### Measurement of serum CK, serum LDH and tissue SOD

Blood samples were collected at 24 hours after reperfusion and plasma was isolated. The activity of creatine kinase (CK) and lactate dehydrogenase (LDH) in plasma were measured using creatine kinase activity assay Kit (Sigma) and LDH activity assay kit (sigma) according to the manufacturer’s instructions. Ventricles were crushed to a powder using liquid nitrogen and homogenized in saline with the weight/volume ratio of 1:10. After centrifuging for 10minutes at 3,500 rpm, the supernatants were withdrawn for SOD activity measurement using SOD assay kit-WST (Dojindo) according to the manufacturer’s instructions. Bradford protein assay was performed to determine the protein concentration.

### Histology assay

Histological studies were performed as previously described^44^. Briefly, animals were sacrificed and the hearts were perfused with 20% KCl. After being fixed with zinc fixative solution (BD Pharmingen) and dehydrated by alcohol, the samples were embedded by paraffin and sectioned into 5 μm slides. The sections were processed for immunostaining, including Masson’s trichrome, immunofluorescence and TUNEL assay (in situ cell death detection kit, Roche). Images were captured by Aperio (Leica Biosystems, Buffalo Grove, IL, USA) and a confocal microscope (Nikon, Melville, NY, USA) and analyzed by Image J software.

### Isolation of neonatal ventricular myocytes (NRVM) and simulated ischemia reperfusion(sI/R) in vitro

NRVM were isolated from postnatal day 1-3 SD rats as previously described^45^. Briefly, neonatal rat hearts were minced into small pieces and transferred into conical tube. The tissues were digested in Trypsin-EDTA (0.25%) at 4°C overnight, then subjected to 1mg/mL type II collagenase at 37°C for 30 minutes. The cell suspension was collected and then centrifuged at 1000rpm for 5 minutes. The resultant pellet was resuspended, and the neonatal cardiac fibroblasts were removed by 2 repeats of 45 minutes plating at cell incubator. The enriched NRVM were then cultured in 5% horse serum for 2 days then changed into serum free medium for further examination. For sI/R, cells were cultured in simulated ischemia medium^16^ and subjected to hypoxia in a chamber with 94% N_2_, 1% O_2_, 5% CO_2_. After 2 hours of hypoxia, the cells were then reperfused in DMEM for 4 hours in serum free medium at 95% air and 5% CO_2_.

### Measurement of Acetyl-CoA

Acetyl-CoA was measured using acetyl-CoA assay kit (Biovision) according to the manufacturer’s instructions. For tissues, hearts were weighted and pulverized, then subjected to 400 μL of 1M perchloric acid/100 mg tissue. For cell culture, 5 million NRVMs were isolated and lysed in RIPA buffer. The lysate was deproteinized by 1M perchloric acid. The deproteinized supernatant was neutralized by 3M KHCO_3_. The supernatant was then measured acetyl-CoA following the standard kit protocol. The size of NRVM was estimated at 6000μm^3^ and the concentration of acetyl-CoA in cells was calculated accordingly.

### Western blot

Proteins were extracted in lysis buffer followed by centrifugation at 4°C for 15 minutes at 12,000 rpm. Protein concentration was measured by Bradford protein assay and 40 μg of total protein was separated by SDS-PAGE and then transferred to PVDF membranes. The membranes were blocked with 5% nonfat dry milk for 1 hour at room temperature and then incubated with primary antibodies overnight at 4°C. After 3 washings with TBST, the membranes were incubated with secondary antibody in TBST solution for 1 hour at room temperature. After 3 washings, the membranes were scanned and quantified by Odyssey CLx Imaging System (LI-COR Biosciences, USA).

### RNA-seq

Total RNA was extracted from the LV tissues using Trizol following manufacture’s protocol. The total RNA was treated with DNAse Turbo to remove genomic DNA. RNA quality was assessed using Agilent Bioanalyzer Nano RNA Chip. 1 μg of total RNA (RIN > 8) was used to prepare the sequencing library using NEBNext Stranded RNA Kit with mRNA selection module. The library was sequenced on illumina HiSeq 4000 (single end, 50 base pair) at the Sequencing Core of University of Michigan. The RNA-seq data have been deposited in Gene Expression Omnibus with the accession code GSE132515.

### RNA-seq data analysis

RNA-seq data was quantified using Kallisto (Version 0.43.0)^46^ with parameters: --single -b 100 -I 200-s 20 using the Rnor6.0 (ensembl v91). The estimated transcript counts were exported by tximport^47^ for Deseq2 analysis^48^ Differential expression was then calculated using Deseq2 default setting. Gene Ontology analysis was performed using GSEA^23^.

### DHE staining

For in vitro staining, 5mM of freshly prepared DHE solution was directly added to medium to final concentration of 5μM and cultured at 37°C for 30 minutes. The cells were washed 3 times of PBS and observed under microscope or dissociated for FACS analysis.

For in vivo staining, heart tissues were embedded in OCT immediately after harvested. Tissues were then sectioned at 20 μm. 5μM of DHE solution was directly apply to sections for 30 minutes at 37°C. After 3 washes of PBS, the sections were mounted and observed under the microscope.

### ChIP

ChIP experiments were performed as previously described^49^. Cells were fixed in 1% formaldehyde and quenched with 0.125M glycine. Nuclei pellets were then harvested and digested with MNase at 37°C for 2 minutes. After brief sonication, chromatin solution was incubated with Dynabeads and antibodies against H3K9ac or pan H3 control overnight. Beads were washed four times with LiCl wash buffer and one wash of TE buffer then eluted with elution buffer at 65°C. DNA was purified using Bioneer PCR purification kit. Enrichment of immunoprecipitated DNA was normalized to pan H3 by quantitative PCR.

### Statistical analysis

GraphPad Prism Software (version 7) was used for statistical analysis. Data were expressed as the mean± SD. Statistical comparisons between two groups were performed by Student’s t test, and more than two groups were performed by two-way or one-way ANOVA followed by post-hoc Tukey comparison. Groups were considered significantly different at *p* < 0.05.

## Acknowledgement

This work was supported by National Institutes of Health (NIH) of United States (1R01HL139735), an Inaugural Grant from the Frankel Cardiovascular Center, an MCube Grant from University of Michigan, and a Pilot Grant from the University of Michigan Health System - Peking University Health Sciences Center Joint Institute for Clinical and Translational Research.

We thank Frankel Cardiovascular Center Physiology and Phenotyping Core for performing all echocardiography examination.

## Author contribution

I.L., S.T., W.G. and Z. W. conceived and performed the experiment and wrote the manuscript. L.L. and Y.G. performed experiments. All authors have read and approved the final version of the manuscript.

## Declaration of Interests

The authors declare no conflict of interests.

## SUPPLEMENTAL MATERIAL

### Supplemental Figure legends

**Supplemental Figure 1.**
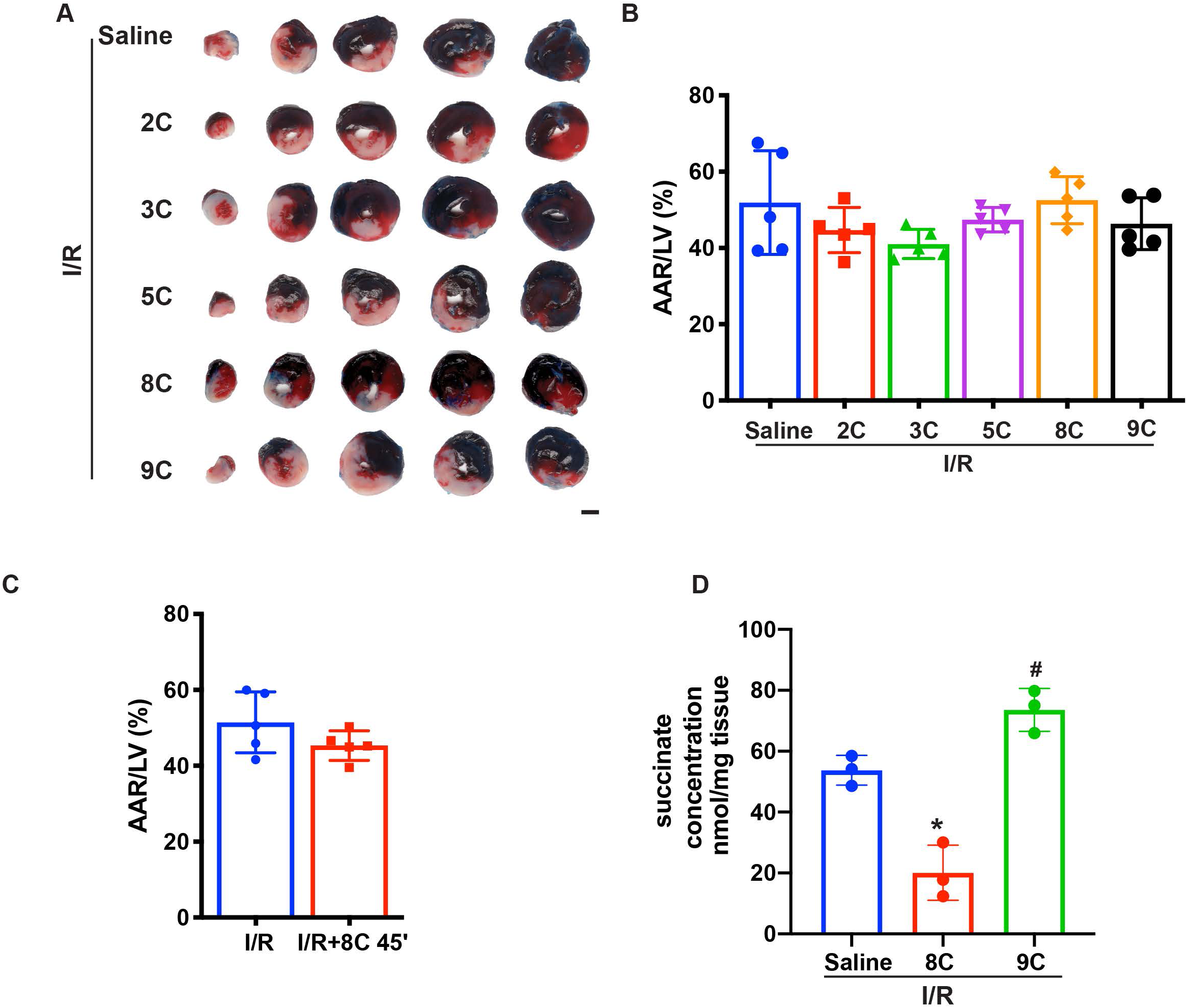
(A) Representative figures of heart sections at 24 hours after I/R in presence of different metabolites. Scale bar: 2.5 mm. (B) Quantification of AAR/LV in Figure S1A by Image J. (C) Quantification of AAR/LV ratio after I/R with or without 8C administration at 45 minutes after ligation. (D) Quantification of Succinate levels in rat hearts after I/R with saline, 8C or 9C treatment. n=3, *P < 0.05, **P < 0.01 vs I/R+saline group; ^##^P < 0.01 vs I/R+8C group.

**Supplemental Figure 2.**
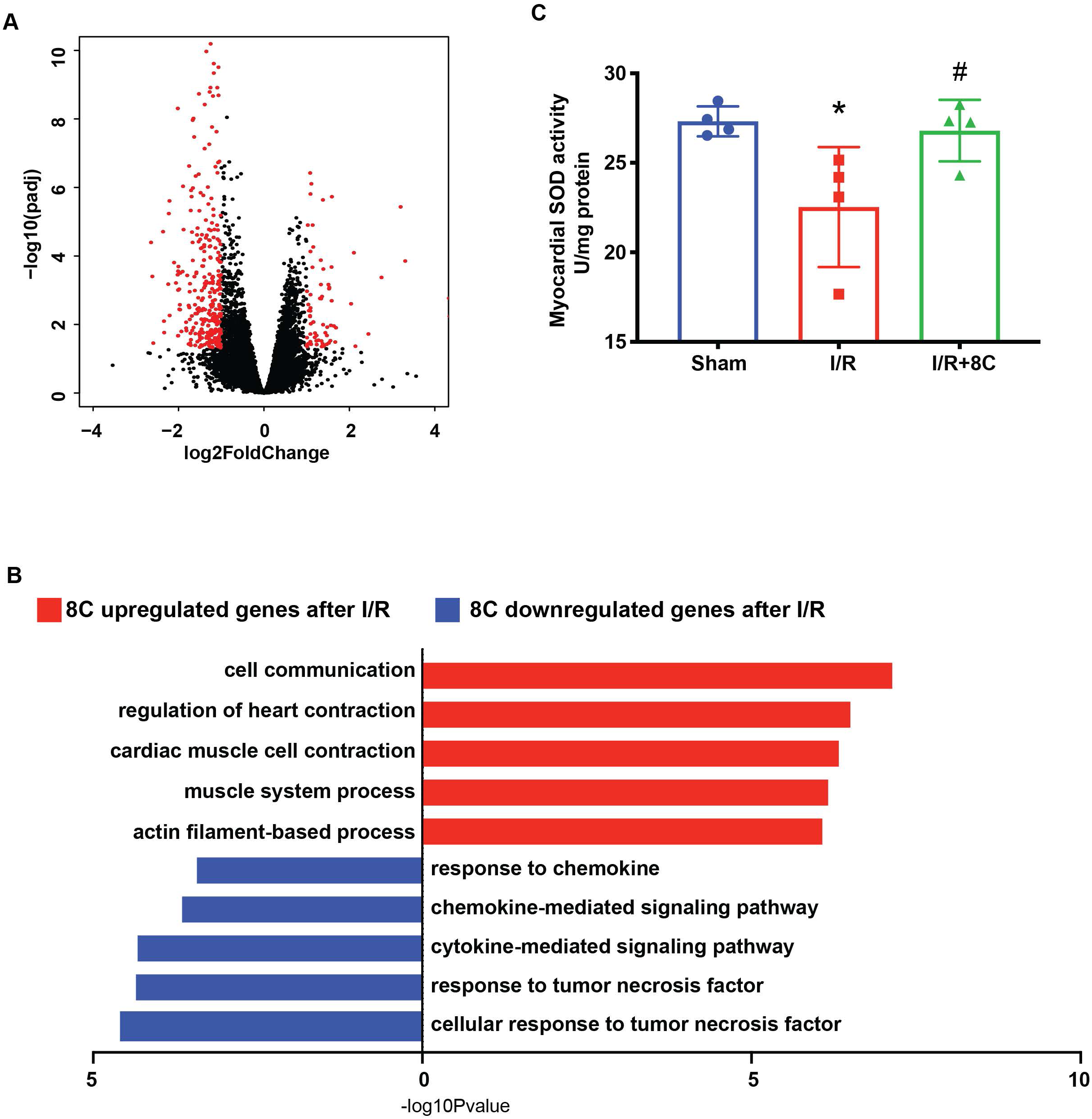
8C administration altered the gene expression after I/R injury. (A) Volcano plot of differential expressed genes in 8C treated and saline treated hearts after I/R. Differential expressed genes with abs(Log2Foldchange) >1, and padj < 0.05 were labelled in red. (B) Top gene set enrichment of differential expressed genes using GO biological process gene sets. (C) Quantification of myocardial SOD activity after I/R with or without I/R. n=3, *P < 0.05 vs Sham group; ^#^P < 0.05 vs I/R group.

**Supplemental Figure 3.**
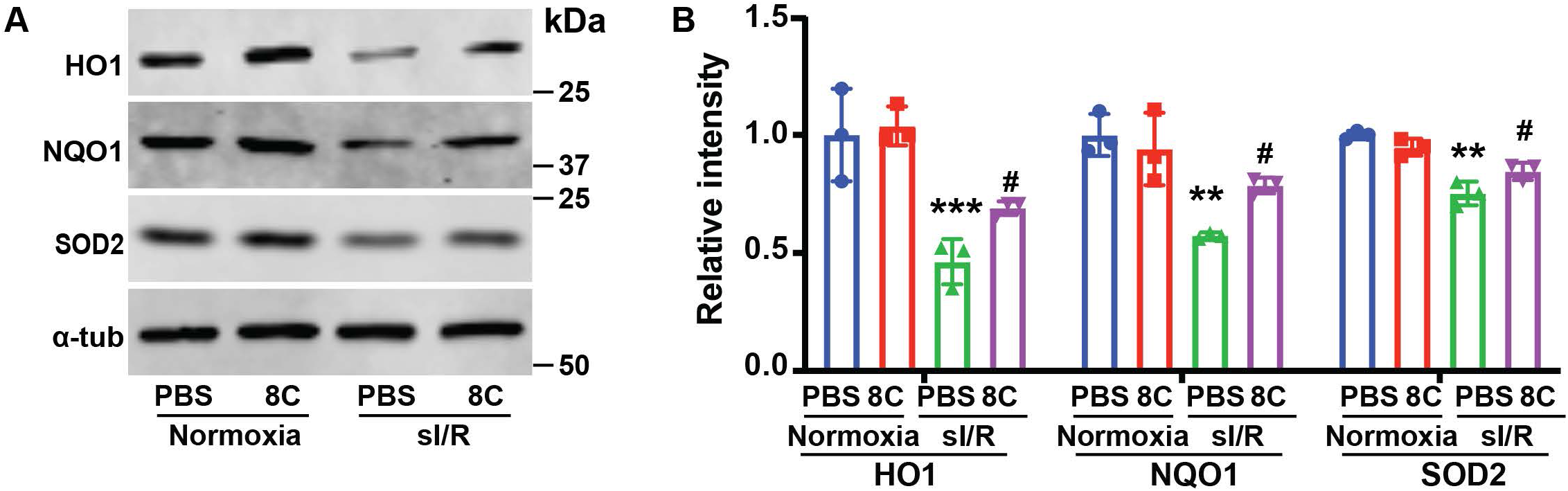
Expression of antioxidant genes with 8C treatment after sI/R. (A) Western blot and (B) quantifications of HO1, NQO1, and SOD2 in NRVM after sI/R. n=3, *p<0.05, **p<0.01, ***p<0.001, vs Normoxia+PBS; ^#^p<0.05, ^##^p<0.01 vs sI/R+PBS.

**Supplemental Figure 4.**
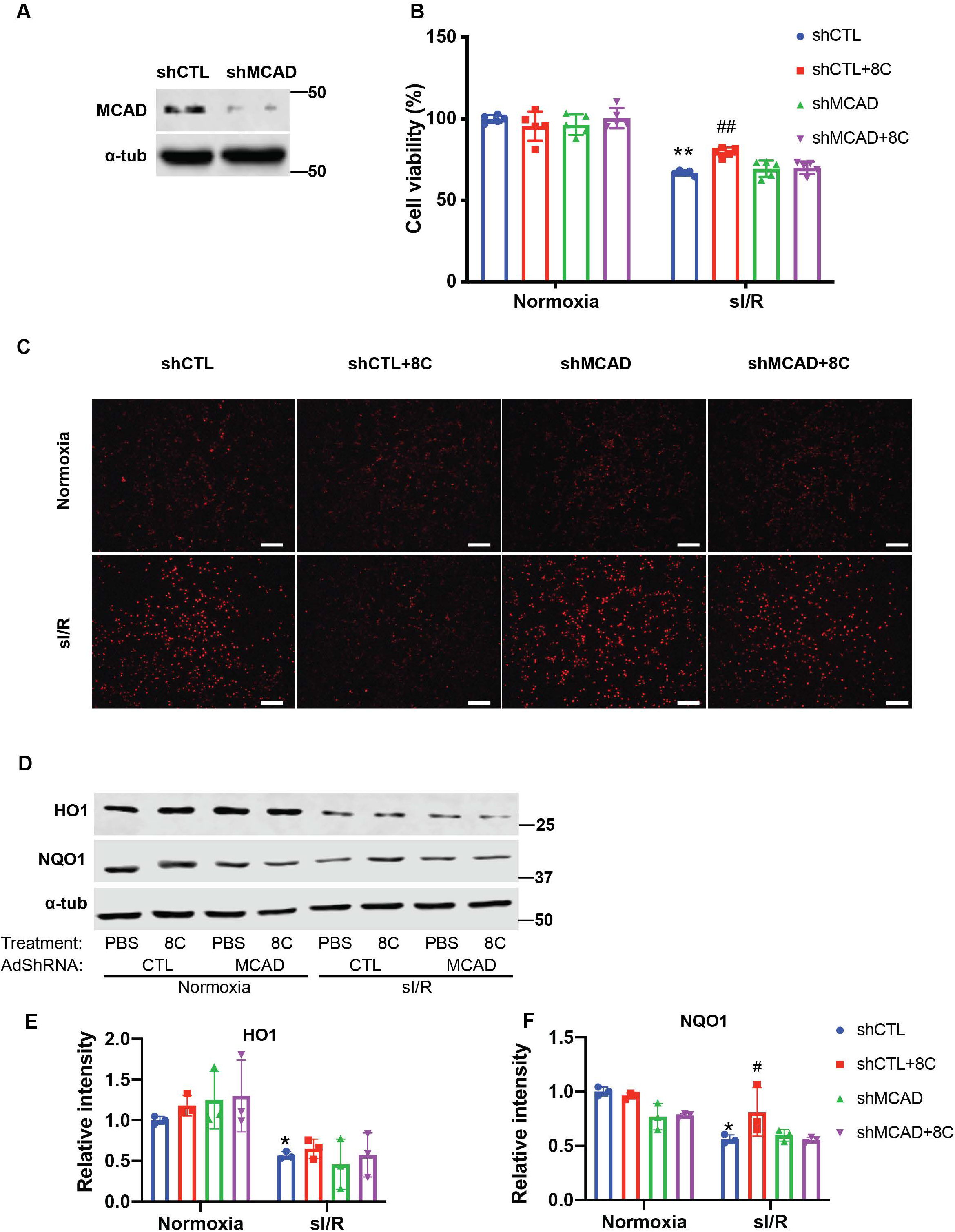
Knockdown of MCAD is required for alleviating ROS accumulation. (A) Western blot showed knockdown of MCAD by adenovirus shRNA. (B) Cell viability measurement in NRVM at shown condition using CCK8 detection kit. (C) NRVM cellular ROS levels are indicated by DHE staining after sI/R treatment. Scale bar: 200μm. (D-F) Western blot and quantifications of HO1 and NQO1 in NRVM after sI/R. n=3, *p<0.05, **p<0.01, vs Normoxia+PBS+shCTL; ^#^p<0.05, ^##^p<0.01 vs sI/R+PBS+shCTL.

**Supplemental Figure 5.**
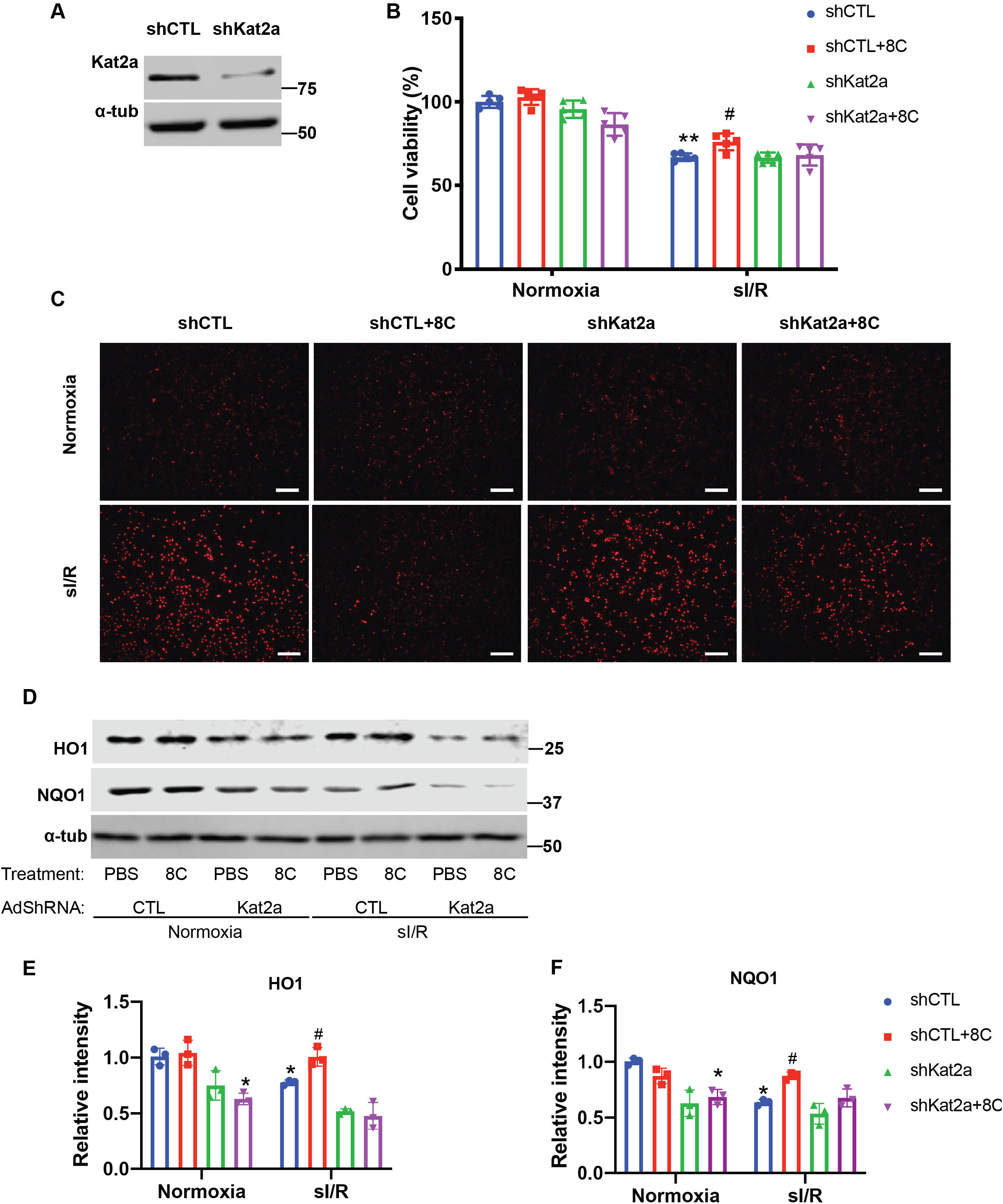
Knockdown of Kat2a is required for alleviating ROS accumulation. (A) Western blot showed knockdown of Kat2a by adenovirus sh-RNA. (B) Cell viability measurement in NRVM at indicated condition using CCK8 detection kit. (C) NRVM cellular ROS levels are indicated by DHE staining after sI/R treatment. Scale bar: 200μm. (D-F) Western blot and quantifications of HO1 and NQO1 in NRVM after sI/R. n=3, *p<0.05, **p<0.01, vs Normoxia+PBS+shCTL; ^#^p<0.05 vs sI/R+PBS+shCTL.

**Supplemental Figure 6.**
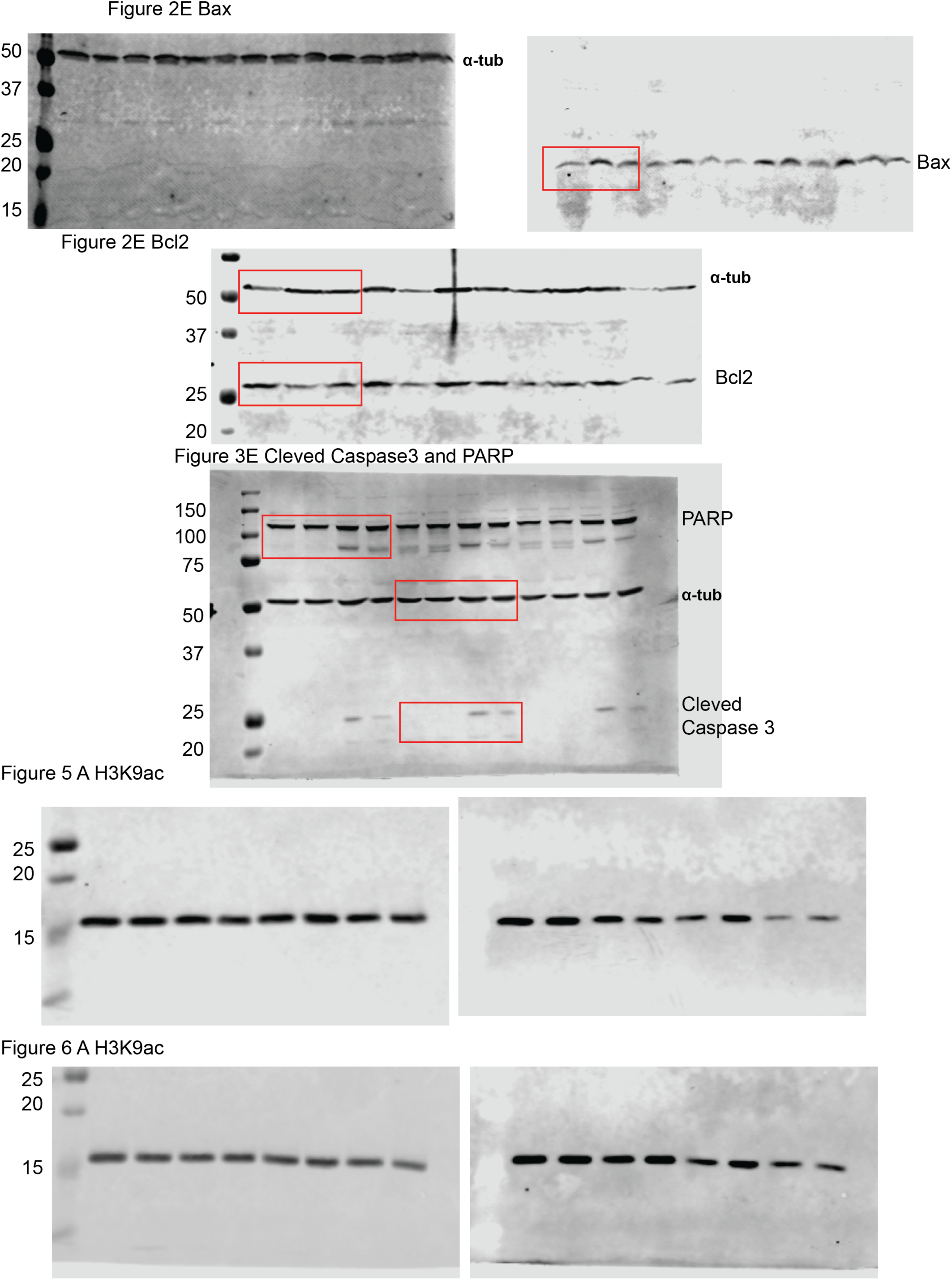
Complete western blot images before cropping into result figures.

**Supplemental Figure 7.**
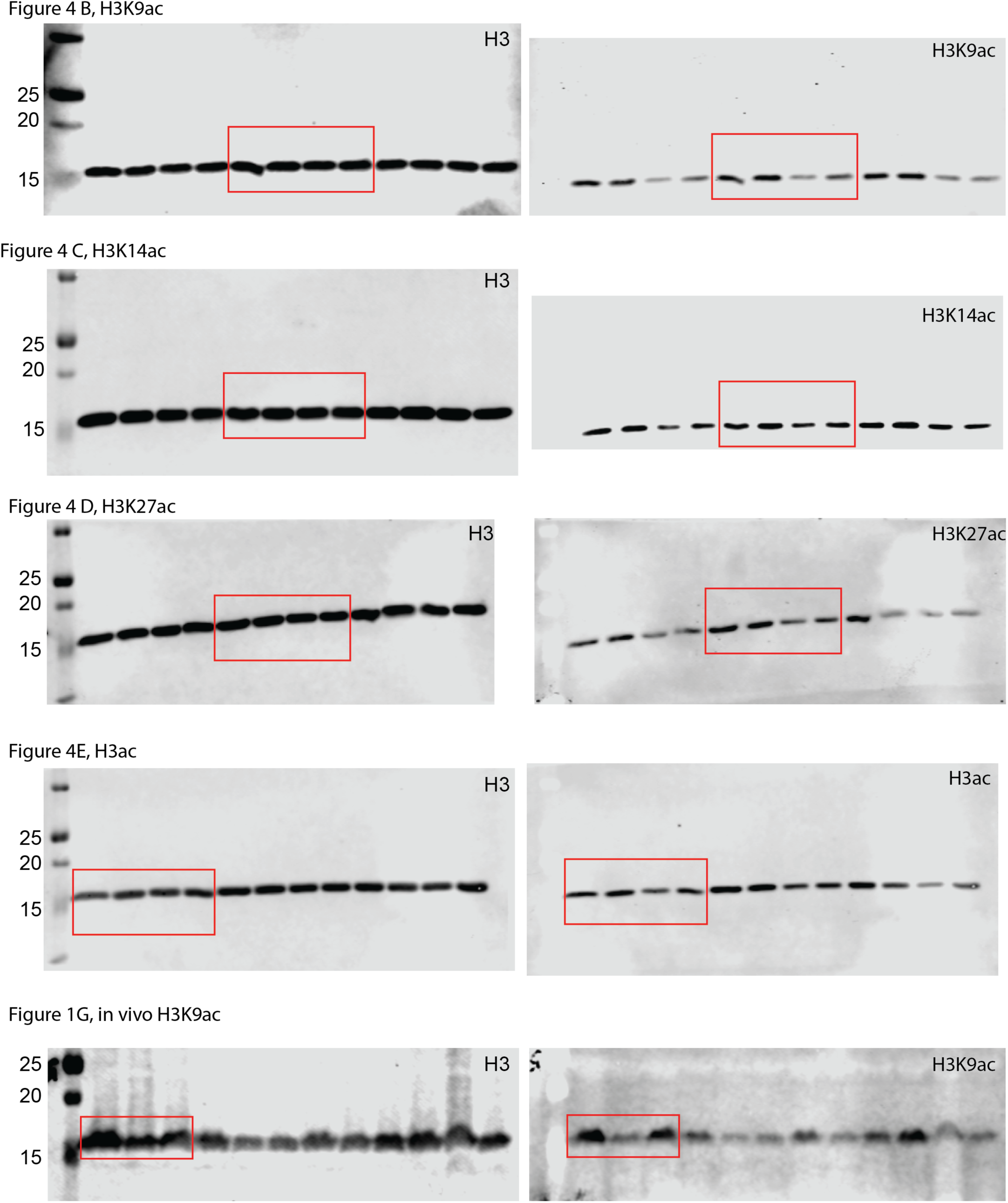
Complete western blot images before cropping into result figures.

**Supplemental Figure 8.**
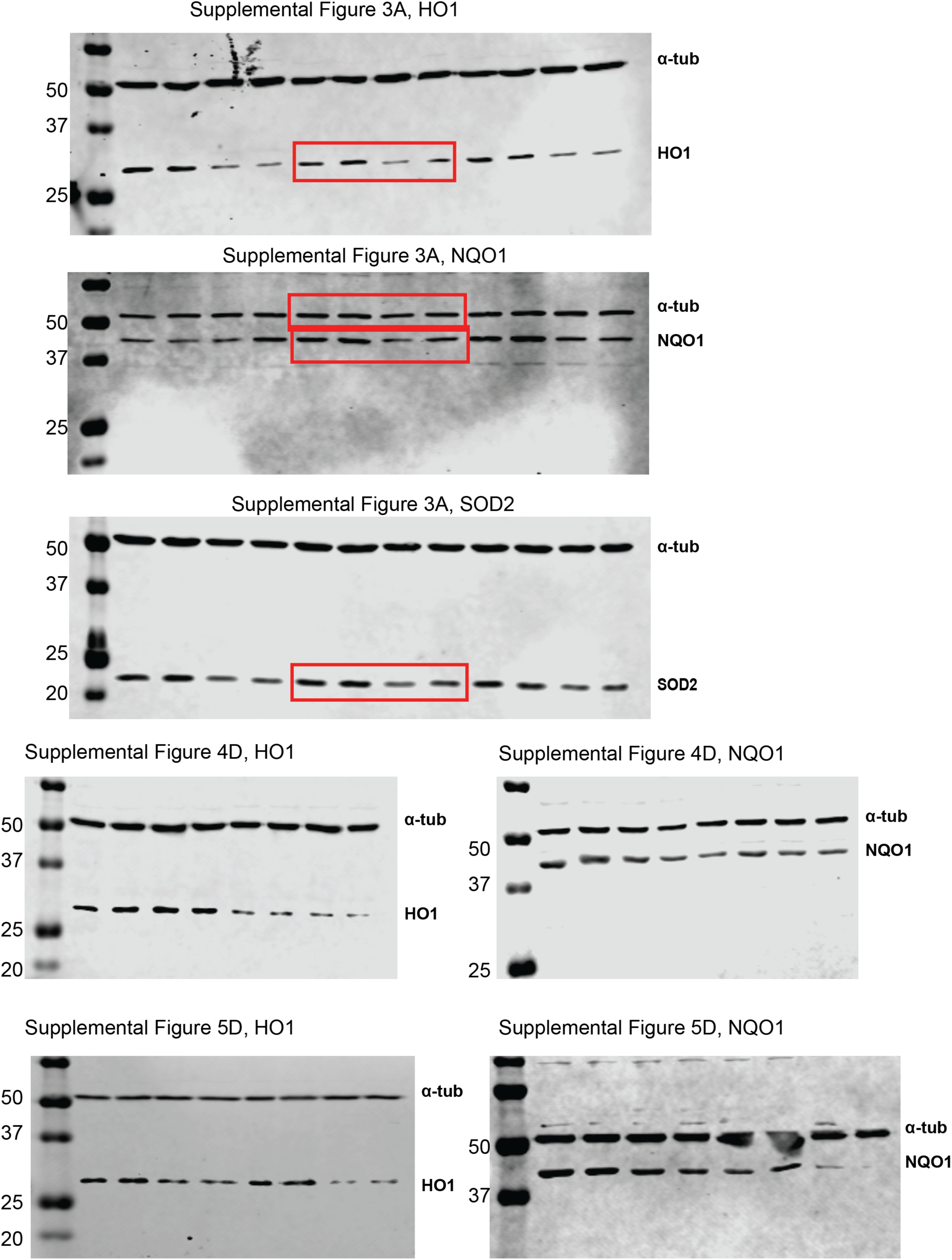
Complete western blot images before cropping into result figures.

